# Consensus Development of a Modern Ontology of Emergency Department Presenting Problems – the HierArchical Presenting Problem ontologY (HaPPy)

**DOI:** 10.1101/126870

**Authors:** Steven Horng, Nathaniel R. Greenbaum, Larry A. Nathanson, James C McClay, Foster R. Goss, Jeffrey A. Nielson

**Affiliations:** Beth Israel Deaconess Medical Center — Harvard Medical School — Department of Emergency Medicine Boston, MA, USA; University of Nebraska Medical Center College of Medicine — Department of Emergency Medicine, Omaha, NE, USA; University of Colorado Hospital — University of Colorado School of Medicine - Department of Emergency Medicine, Aurora, CO, USA; Northeastern Ohio Medical University — University Hospitals Samaritan Medical Center, Ashland, OH, USA

**Keywords:** chief complaint, presenting problem, Emergency Department, Emergency Medicine, ontology

## Abstract

**Objective:** Numerous attempts have been made to create a standardized ‘presenting problem’ or ‘chief complaint’ list to characterize the nature of an Emergency Department visit. Previous attempts have failed to gain widespread adoption as none were freely sharable and contained the right level of specificity, structure, and clinical relevance to gain acceptance by the larger emergency medicine community. Using real-world data, we constructed a presenting problem list that addresses these challenges.

**Materials and Methods:** We prospectively captured the presenting problems for 180,424 consecutive emergency department patient visits at an urban, academic, Level I trauma center in the Boston metro area. No patients were excluded. We used a consensus process to iteratively derive our system using real-world data. We used the first 70% of consecutive visits to derive our ontology; followed by a 6 month washout period, and the remaining 30% for validation. All concepts were mapped to SNOMED-CT.

**Results:** Our system consists of a polyhierarchical ontology containing 692 unique concepts, 2,118 synonyms, and 30,613 non-visible descriptions to correct misspellings and non-standard terminology. Our ontology successfully captured structured data for 95.9% of visits in our validation dataset.

**Discussion and Conclusion:** We present the HierArchical Presenting Problem ontologY (HaPPy). This ontology was empirically derived then iteratively validated by an expert consensus panel. HaPPy contains 692 presenting problem concepts, each concept being mapped to SNOMED-CT. This freely sharable ontology can help to facilitate presenting problem based quality metrics, research, and patient care.

## 1. BACKGROUND AND SIGNIFICANCE

The precipitating reason for a visit is an important data element that is captured when a patient presents to an emergency department (ED) and is often one of the first questions providers ask a patient. This information is used to guide the patient’s initial clinical care, and when aggregated, serves as a valuable tool for understanding patterns of patient visits for administrative and research purposes. Often referred to as a *chief complaint* or *reason for visit,* there is no formalized definition or required vocabulary for recording this information in emergency department information systems.

The high degree of variability in recording this information both within and between hospitals greatly hampers reuse of this information for clinical, research, or quality purposes. For example, a specific set of symptoms can be recorded by different providers as “chest pain”, “CP” or “cardiac pain”, making evaluation of data difficult. This is further complicated by the prevalence of unstructured free-text entries, which commonly have misspellings, local abbreviations and other errors that are unsuitable for use in computerized decision support and for secondary analysis.^1,2^ The need for standardization has been well described in the past^3^, but no solution has received wide adoption.

Over the last 15 years attempts have been made to create a standardized method of recording chief complaints^4^. Prior attempts have failed to gain widespread acceptance for various reasons: they may not be freely sharable or may not have had the right level of specificity, structure, and clinical relevance to gain acceptance by the larger emergency medicine community^5^. Emergency department information system (EDIS) vendors and other commercial vendors^6^ offer vocabularies, but they cannot be used to compare data across EDs who do not have access to these proprietary vocabularies. Other efforts at non-proprietary vocabularies^7,8^ were developed, but lacked the granularity for effective secondary use. Attempts to solve this by computerized natural language processing did not yield sufficient sensitivity and specificity.^9^ Furthermore, most work in the field is in the secondary use of chief complaints to classify visits for syndromic surveillance^10–18^, rather than characterizing a presenting problem for an emergency department visit.

To address these issues, the American College of Emergency Physicians Section of Emergency Medicine Informatics was tasked with developing a freely available, standardized vocabulary for the “presenting problem” (PP) suitable for use in any ED electronic health record (EHR). We make a distinction between the “chief complaint” (CC), defined as the patient’s own words^19^ and the PP which is a provider’s clinical interpretation of the patient’s concerns.

When patients present to the emergency department (ED), they share a reason for their visit with the initial provider. These first spoken words are the chief complaint (CC). While classically taught in medical school that the CC should be in the patient’s own words, there has been a shift towards recording the CC using a list of standardized terms recognized by the local electronic health record (EHR). The categorized version of the complaint requires a transformation from a patient’s view of the problem to the provider’s interpretation of the problem. Unfortunately, this transformed term is still often referred to as “chief complaint”, even though it is a distinct new entity. We choose to use the term chief complaint as originally intended, and term the new entity the “presenting problem” (PP). The PP is the provider interpretation of the patient’s chief complaint (**Table 1**).We collect the PP (provider’s perspective) in a text field, separate from other free-text. We do not collect a separate chief complaint (patient’s perspective), although some EHR’s might.

**Table 1:**
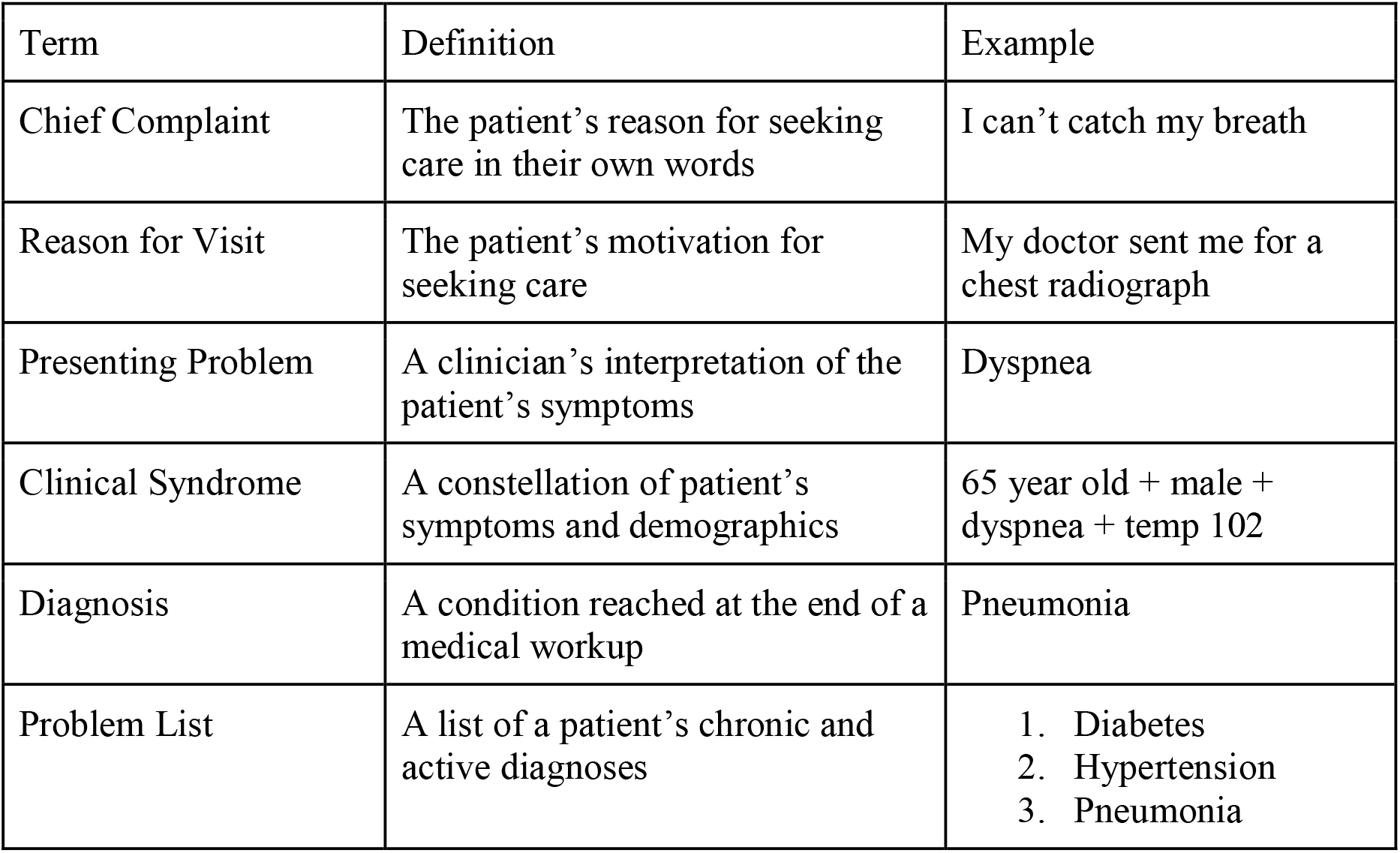
Definition of Terms.

| Term | Definition | Example |
| --- | --- | --- |
| Chief Complaint | The patient's reason for seeking care in their own words | I can't catch my breath |
| Reason for Visit | The patient's motivation for seeking care | My doctor sent me for a chest radiograph |
| Presenting Problem | A clinician's interpretation of the patient's symptoms | Dyspnea |
| Clinical Syndrome | A constellation of patient's symptoms and demographics | 65 year old + male + dyspnea + temp 102 |
| Diagnosis | A condition reached at the end of a medical workup | Pneumonia |
| Problem List | A list of a patient's chronic and active diagnoses | 1. Diabetes<br>2. Hypertension<br>3. Pneumonia |

In 2006, a group of 40 stakeholders held a national consensus meeting to develop the framework for a standardized chief complaint vocabulary^5^; a ten-point consensus plan was conceived. We executed portions of the plan to construct an ontology of presenting problems that can be used by a wide variety of users, leveraging existing standards, and ready for external validation studies.

An ontology is a controlled vocabulary of well-defined terms with specified relationships between those terms, capable of interpretation by both humans and computers.^20^ This structure is known as an “ontology” and means that concepts such as “broken tooth” and “dental abscess” are understood by the computer to be related to each other as a “tooth disorder” concept.

We convened a multi-center physician and nurse expert panel of emergency informatics experts with a goal of creating a PP ontology that fosters clinical decision support, standardized clinical pathways^21^, quality measures^22, 23^, and research. It should also support information exchange from prehospital providers^24^ as well as with the public health system for syndromic surveillance. Leveraging informatics techniques, the end product is more than just a simple list of words — it also includes the relationships between terms, creating a powerful way to easily group and categorize related visits.

## 2. OBJECTIVES

Our goal was to develop a standardized presenting problem ontology that would be precise and easy to use while capturing at least 95% of all presenting problems at the primary institution site.

## 3. METHODS

### 3.1 Overview

We conducted a quality improvement project over a 3 year period to develop and validate a standardized emergency medicine presenting problem ontology. We call this the HierArchical Presenting Problem ontologY (HaPPy). We collected free text presenting problems as well as patient demographics.

### 3.2 Goals of the Investigation

The following goals motivated the development of this ontology:

**Goal 1 - Standardize the collection of Emergency Department Presenting Problems**

**Goal 2 - Develop an ontology that can be easily deployed to existing EHR’s**

**Goal 3 - Reliably cohort patients for quality measurement, specifically for development of electronic clinical quality meaures (eCQM’s)^25^**

**Goal 4 - Trigger presenting problem-based decision support**

**Goal 5 - Reliably cohort patients for research and quality improvement**

### 3.3 Setting and Selection of Participants

The study was performed in a 55,000 visits/year Level I trauma center and tertiary, academic, adults-only, teaching hospital. All consecutive ED patient visits between 3/10/2013 and 5/29/16 were included in the study. The first 106,695 consecutive visits (70%) were used to derive the ontology, followed by a 6 month washout period, and finally a validation period of 45,687 (30%) patient visits. No visits were excluded. The EHR used was custom developed at the institution.

### 3.4 Iterative Presenting Problem Development

Our grounded theory approach builds upon SNOMED CT, an internationally developed and maintained hierarchical ontology, by using real-world data captured at the point of care. It improves over previous list-based approaches and provides a foundation for future PP advances. We employed the four usability requirements for structured documentation described by Rosenbloom et al.^26^ to create a system that is easy to use and clinically meaningful.

To create the presenting problem vocabulary, we examined all consecutive presenting problems entered into our EHR from 10 March 2013 to 11 February 2015. After an initial period of data collection, unstructured (free text) presenting problems were reviewed by the committee to create an initial PP dataset. In an iterative process, free text entries that were not already part of our ontology were reviewed to identify candidates for inclusion. During each review, free text presenting problems not yet in the vocabulary were sorted by frequency with the most frequent presenting problems considered first for addition. Lexical variants, synonyms, and misspellings were also captured.

Free text presenting problem terms were normalized and then manually mapped to concepts in SNOMED CT (March 2013) and the US Extension to SNOMED CT (September 2012). The level of granularity, as well as proper mapping to SNOMED CT was performed using the consensus process defined below. The updated presenting problem list, along with its synonyms and lexical variants, was then re-deployed to users. After a brief collection period, this process was repeated until we reached the a priori termination point of 95% coverage. This iterative process is illustrated in **Figure 1**.

**Figure 1:**
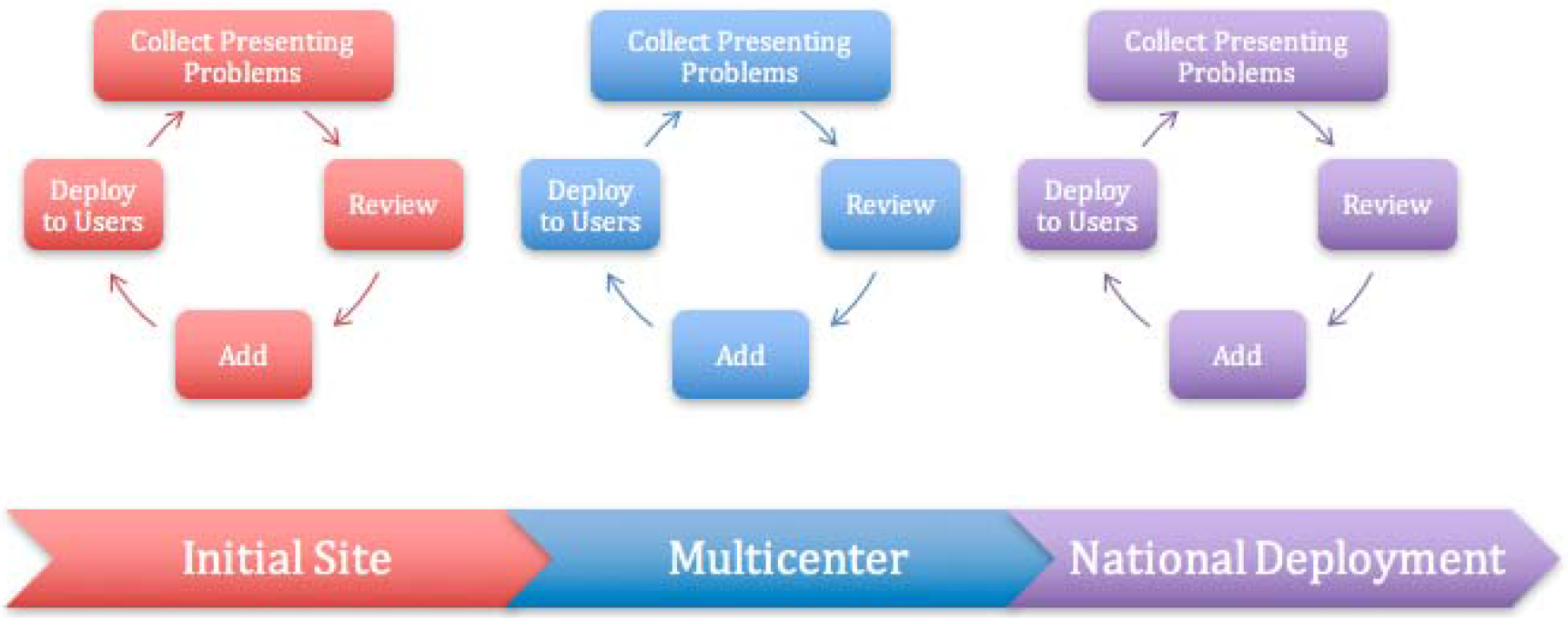
Iterative Presenting Problem Development.

SNOMED CT was used as the foundation vocabulary because it is licensed for use in the United States, freely available, internationally accepted, and regularly maintained and adheres to the principles of a formal ontology. SNOMED provides a structured set of concepts and synonyms, linked via well-defined relationships, allowing for complex computational queries based on these relationships. For example, a search for ‘abdominal pain’ allows the user to query all its child terms (right sided abdominal pain, left lower quadrant abdominal pain, epigastric pain, etc) with a single search. SNOMED CT is also a designated terminology for Meaningful Use regulations and the terminology standard for encoding patient problems in EHRs, making it uniquely suited for development of a PP ontology. Our PP ontology is composed of two parts: 1) a reference set (subset) of the SNOMED CT terminology that represents ED presenting problems and 2) an interface terminology that consists of all the lexical variants end-users can enter into the EHR to express those terms.

In this ontology, we created a subset of SNOMED CT. Although SNOMED CT could have been used in its native, clinical users would have had difficulty in choosing from 399,117 unique concepts. Clinical users are not expert ontologists, and would be unlikely to reliably select from the correct hierarchy. Furthermore, an ontology of 399,117 unique concepts would be difficult for users to learn and use.

### 3.5 Consensus Methodology

The content was developed by a group of 8 emergency physicians and emergency nurses at urban, academic medical centers who also are leaders in clinical informatics. Unanimous consensus was achieved after 9 meetings over 2 years, with a combination of in-person meetings at national conferences and teleconference. The group developed a set of rules to determine which problems should be included in the vocabulary. The group also the PP terms to SNOMED CT. The group also developed a set of heuristics to determine whether a problem should be included in the vocabulary.

**Heuristic 1 - Would the addition of the presenting problem change clinical thinking, workflow, or analysis?**

**Heuristic 2 - Is the level of granularity appropriate for a triage nurse?**

**Heuristic 3 - Is the presenting problem supported by current user behavior?**

**Heuristic 4 - Does the presenting problem reduce ambiguity and improve communication?**

The consensus process also determined how to appropriately map each presenting problem to the SNOMED CT hierarchy as the same concept can be represented in multiple hierarchies within SNOMED. In this study, we represent signs and symptoms in the ‘Clinical Findings’ hierarchy, diagnoses in the ‘Disorders’ hierarchy, and events in the ‘Events’ hierarchy. For example, the concept “abdominal pain” was mapped to “Abdominal Pain (finding)”, which is in the clinical Finding hierarchy with semantic type “Sign or Symptom”. Alternatively, it could also have been mapped to “On examination - abdominal pain (finding)”, also under the clinical findings hierarchy, but with semantic type finding. Since abdominal pain in this context is most likely a patient’s reported symptom of abdominal pain, rather than a triage nurse elicited abdominal pain from a physical exam, it was more appropriate to map this to the former concept. Another alternative would be “Abdominal pain characteristic (observable entity)”. However, in this context, the abdominal pain would be most likely a patient’s reported sign or symptom, rather than the triage nurse observing abdominal pain.

The group used the software package Gephi^27^ to help visualize the is-a relationships between presenting problems. We suggest that users use the included .gexf graph file in the distribution to explore the ontology.

We chose to exclude some PPs that were frequently documented at triage based on the application of the above heuristics by the investigators. For example, the term “lethargy” is used colloquially to describe weakness, fatigue, or sleepiness. However, the clinical definition for a physician is a decreased level of consciousness requiring prompt intervention^28^. This discrepancy between the intent of how lethargy is used, and how it is interpreted, leads to much confusion. Similarly, the presenting problem of “arrest” could be used to indicate a cardiac arrest, a respiratory arrest, or that a patient was detained by law enforcement. Omitting these terms reduces ambiguity and improves communication (Heuristic 4).

Additional omissions include items such as “patient referral for consultation”, “minor complaint”, and “imaging tests”. Although these terms may describe why a patient presented to the emergency department, they do not meaningfully change medical decision making or workflow (Heuristic 1). We, however, did include the concept “transfer”, to denote when a patient was transferred from an outside hospital, as it does meaningfully change workflow. We also built decision support so that transfer could not be used as the only presenting problem.

### 3.6 Interface Terminology

We started our interface terminology by manually reviewing all SNOMED CT descriptions for a particular concept to use as synonyms for a particular concept. We also automatically generated several lexical variants. For example, we generated common abbreviations such as UE for upper extremity, Fx for Fracture, and Lac for Laceration. By proactively generating these terms we increase the likelihood that a user will find the term they are searching for, improving the amount of structured data captured.

We also automatically generated lateralizing prefixes for concepts that we indicated as having laterality. For example, for the concept ‘Arm pain’ our system would automatically generate ‘Left arm pain’, ‘L arm pain’, ‘Lt arm pain’, ‘(L) arm pain’, as well as all the permutations with and without the flanking word, ‘-side’, ‘sided’, and ‘-sided’. Laterality was represented in the codes using post-coordination by adding -R, -L, or -B to the end of the concept. This allows developers to easily recover the base concept, as well as the laterality without having to perform a dictionary lookup (**Table 2**). Though this may appear that we are creating a new pre-coordinated term, we mean this to be post-coordination only, and use a -R, -L, -B for usability by EHR’s only, as most EHR’s are not capable of post-coordination.

**Table 2:**
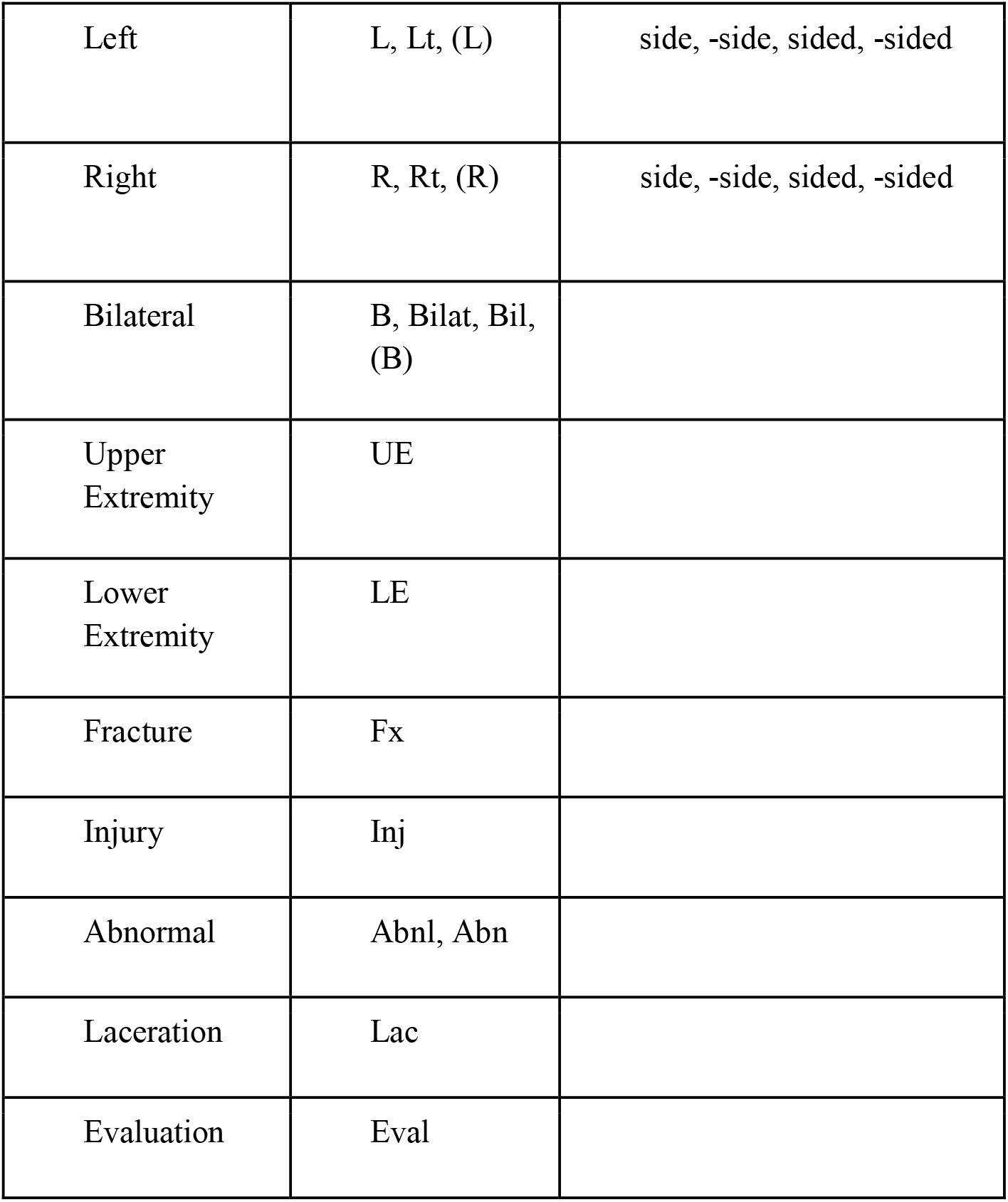
Automated synthesis of lexical variants.

In order to ensure that all anatomic variations for a PP were included, we developed a list of anatomic body parts from all existing PP. We then applied this list of anatomic body parts to existing PP’s to uncover potential PP’s for inclusion. We manually reviewed this potential list of PP for inclusion and included PP according to the heuristics described above. We repeated this process until no additional terms were discovered.

### 3.7 Data Analysis

We derived the ontology using the first 106,695 consecutive visits (70%), followed by a 6 month washout period, and finally a validation set of 45,687 (30%) patient visits.

One or more presenting problems can be documented for each patient visit in our EHR. We defined the primary outcome measure as positive if all of the documented presenting problems listed for the patient were an exact match to a term in our interface terminology.

If any of the presenting problems were not coded (i.e., the triage nurse used free text), we considered the outcome to be negative. For example, a presenting problem entered as “Facial injury / Stab wound to the face” would be recorded as negative since “Facial injury” is in our ontology but “Stab wound to the face” is not. This all or none approach provides the most conservative estimate of the PP ontology’s performance.

## 4. RESULTS

### 4.1 Characteristics of study subjects

A total of 180,424 patient visits were included in the study. These patient characteristics are reported in (**Table 3**).

**Table 3:**
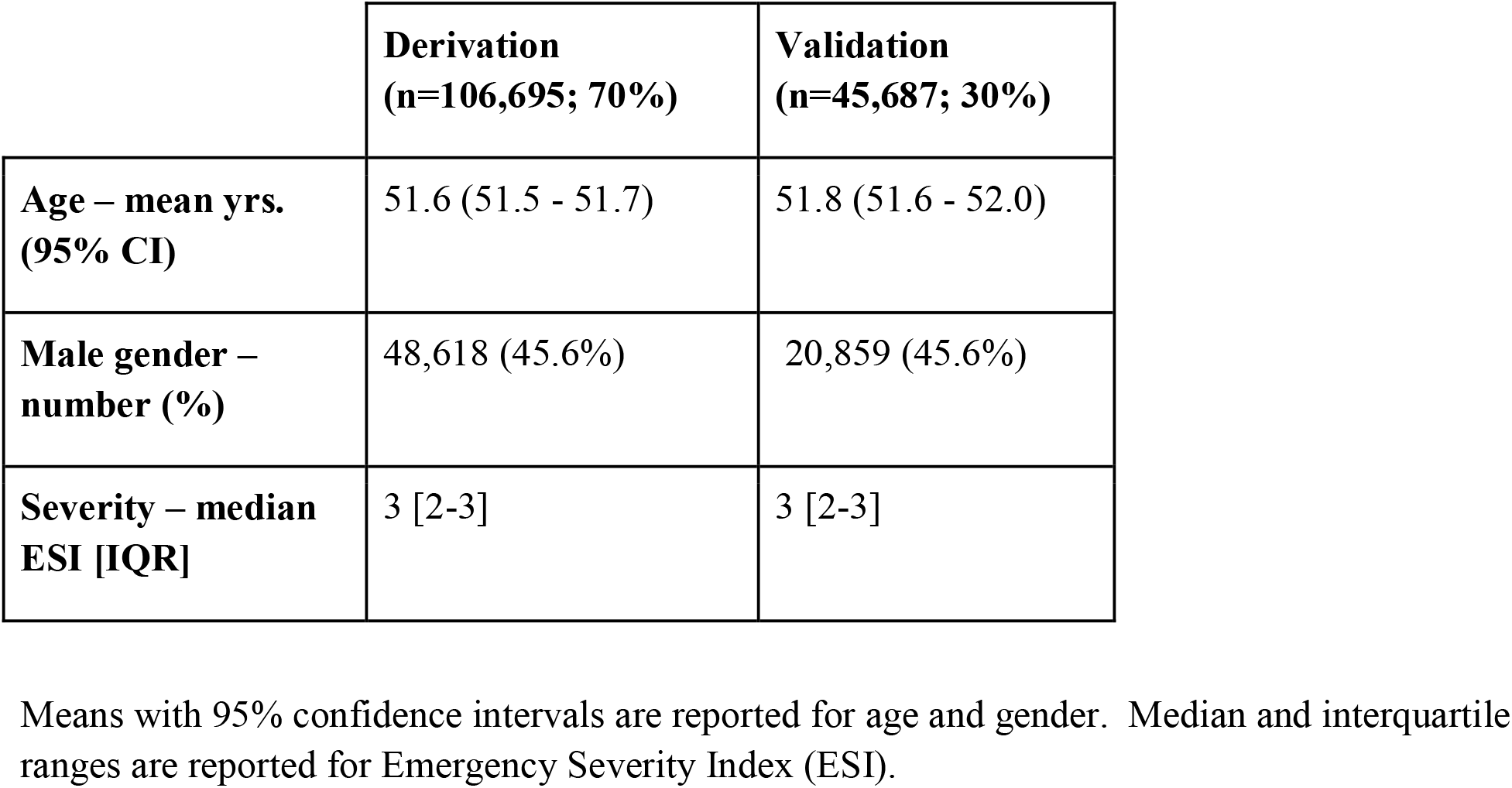
Patient Demographics.

|  | **Derivation**<br>**(n=106,695; 70%)** | **Validation**<br>**(n=45,687; 30%)** |
| --- | --- | --- |
| **Age – mean yrs.**<br>**(95% CI)** | 51.6 (51.5 - 51.7) | 51.8 (51.6 - 52.0) |
| **Male gender –**<br>**number (%)** | 48,618 (45.6%) | 20,859 (45.6%) |
| **Severity – median**<br>**ESI [IQR]** | 3 [2-3] | 3 [2-3] |

Means with 95% confidence intervals are reported for age and gender. Median and interquartile ranges are reported for Emergency Severity Index (ESI).

### 4.2 HierArchical Presenting Problem ontologY (HaPPy)

A total of 692 unique presenting problems were included in our vocabulary. In the validation phase, we found that our presenting problem ontology covered 95.9% of all visits. There were 2,118 synonymous terms that were shown to the user, and an additional 30,614 non-visible descriptions to correct misspellings and non-standard terminology that were not displayed to the user. (**Table 4**)

**Table 4:** Aa Typical Presenting Problem, Synonyms, and Non-visible Descriptions.

| **Presenting Problem [1]** | **Displayed Synonyms [4]** | **Non-visible Descriptions [516]** |
| --- | --- | --- |
| Headache | Headache<br>B Headache<br>L Headache<br>R Headache | H/A<br>HA<br>HEAD ACHE<br>HEADACHE<br>HEADACHES<br>HEAD PAIN<br>MIGRAINE<br>...(more)...<br>(L) HEAD ACHE<br>(L) HEAD PAIN<br>...(more)...<br>RT-SIDED<br>HEADACHES<br>RT-SIDED<br>MIGRAINE |

Relatively few concepts were required to capture the presenting problem for most visits. One-half of all patient visits could be described with just 21 terms, and 75% of visits were described with 58 concepts. Only 121 concepts were required for 90% coverage. For 99.9% coverage, 352 concepts were required, roughly half of our vocabulary. A histogram of presenting problem usage frequency is presented in **Figure 2**. The Top 25 presenting problems are presented in (**Table 5**) and summary of our ontology by SNOMED Semantic Tag appears in (**Table 6**). The top level concepts, ordered by number of children, appears in (**Table 7**).

**Figure 2:**
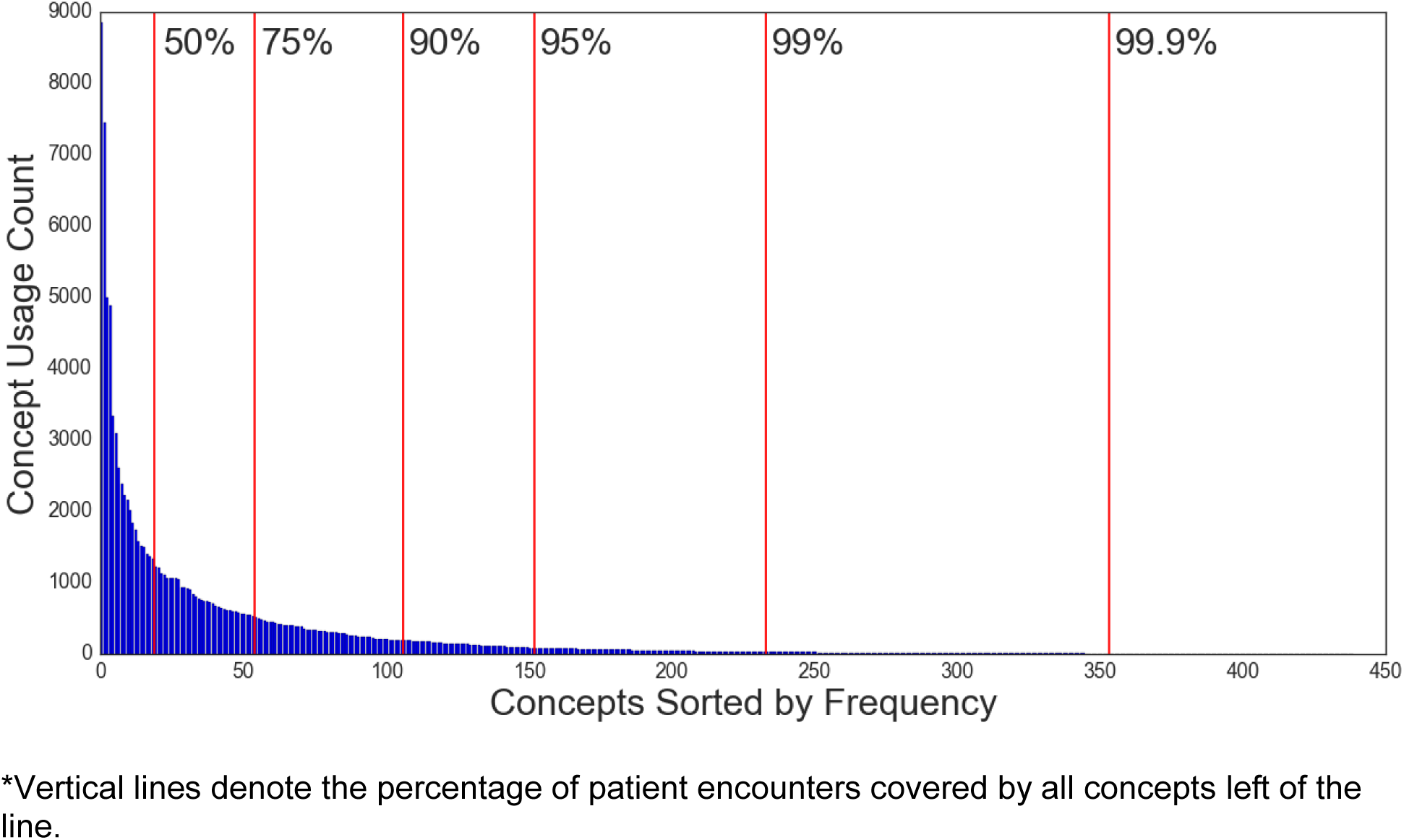
Distribution of Presenting Problem Concept Usage Frequency.

**Table 5:**
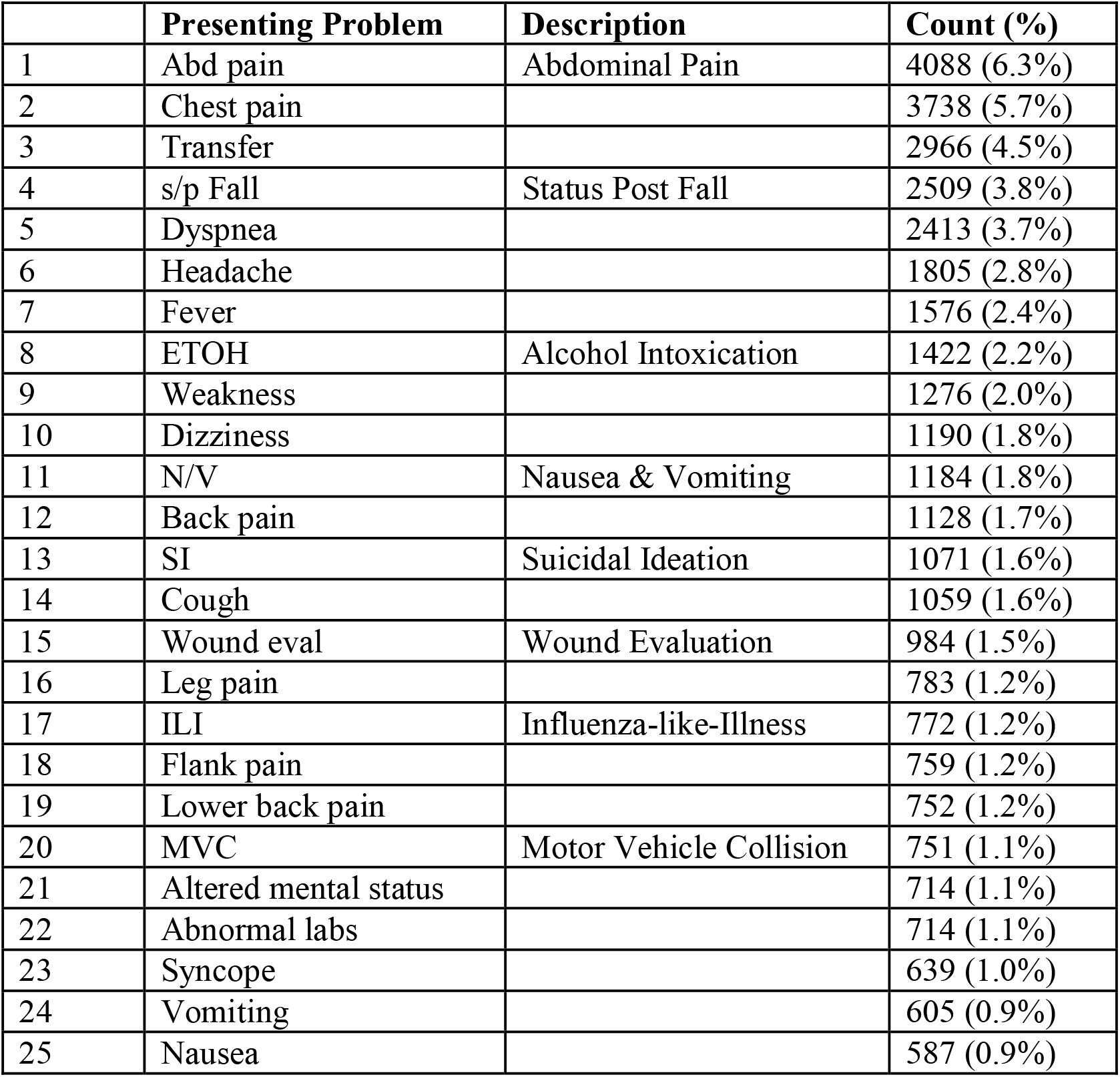
Top 25 Presenting Problems.

|  | **Presenting Problem** | **Description** | **Count (%)** |
| --- | --- | --- | --- |
| 1 | Abd pain | Abdominal Pain | 4088 (6.3%) |
| 2 | Chest pain |  | 3738 (5.7%) |
| 3 | Transfer |  | 2966 (4.5%) |
| 4 | s/p Fall | Status Post Fall | 2509 (3.8%) |
| 5 | Dyspnea |  | 2413 (3.7%) |
| 6 | Headache |  | 1805 (2.8%) |
| 7 | Fever |  | 1576 (2.4%) |
| 8 | ETOH | Alcohol Intoxication | 1422 (2.2%) |
| 9 | Weakness |  | 1276 (2.0%) |
| 10 | Dizziness |  | 1190 (1.8%) |
| 11 | N/V | Nausea & Vomiting | 1184 (1.8%) |
| 12 | Back pain |  | 1128 (1.7%) |
| 13 | SI | Suicidal Ideation | 1071 (1.6%) |
| 14 | Cough |  | 1059 (1.6%) |
| 15 | Wound eval | Wound Evaluation | 984 (1.5%) |
| 16 | Leg pain |  | 783 (1.2%) |
| 17 | ILI | Influenza-like-Illness | 772 (1.2%) |
| 18 | Flank pain |  | 759 (1.2%) |
| 19 | Lower back pain |  | 752 (1.2%) |
| 20 | MVC | Motor Vehicle Collision | 751 (1.1%) |
| 21 | Altered mental status |  | 714 (1.1%) |
| 22 | Abnormal labs |  | 714 (1.1%) |
| 23 | Syncope |  | 639 (1.0%) |
| 24 | Vomiting |  | 605 (0.9%) |
| 25 | Nausea |  | 587 (0.9%) |

**Table 6:**
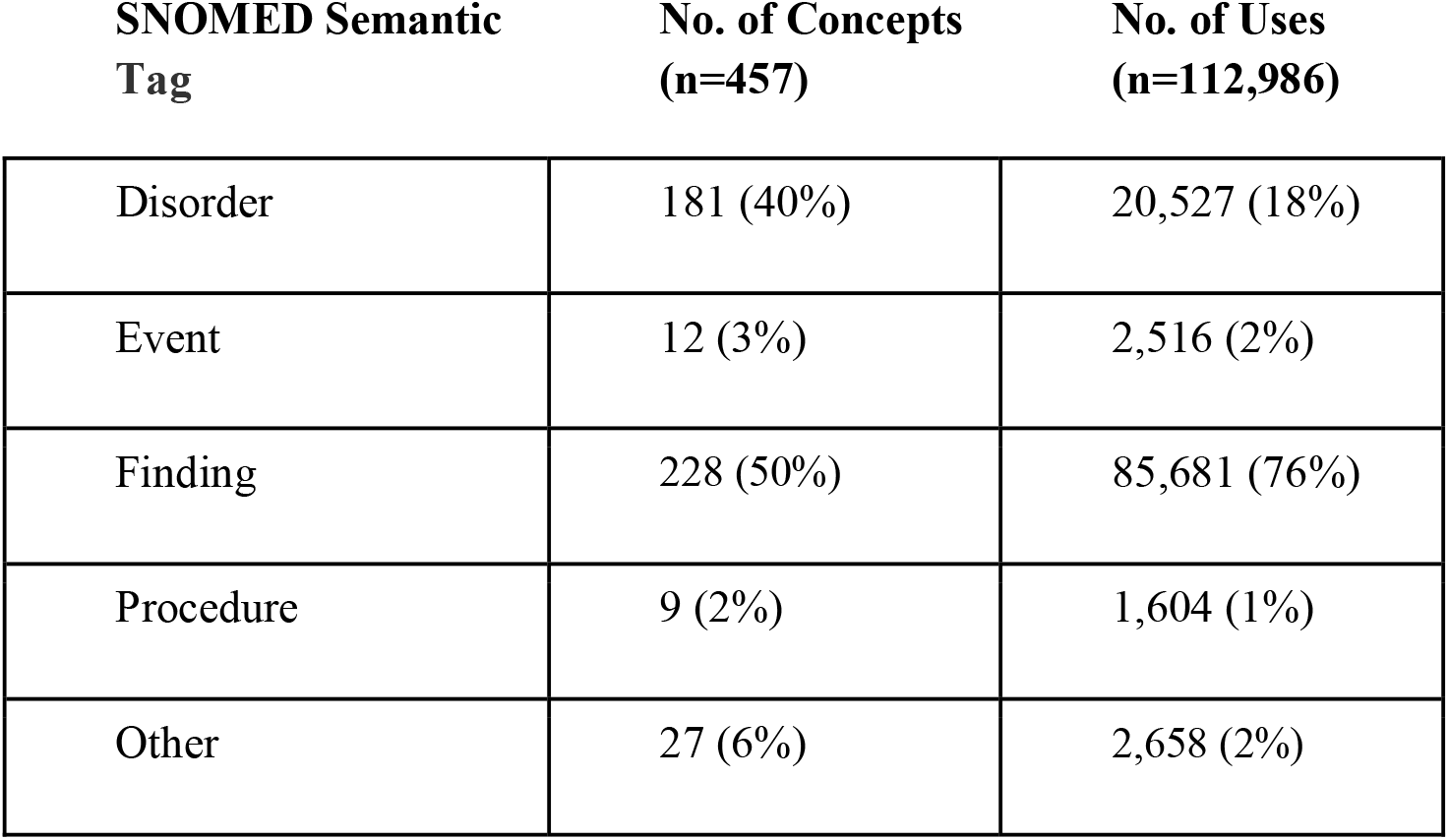
Distribution of Concept Types and Their Usage.

**Table 7:**
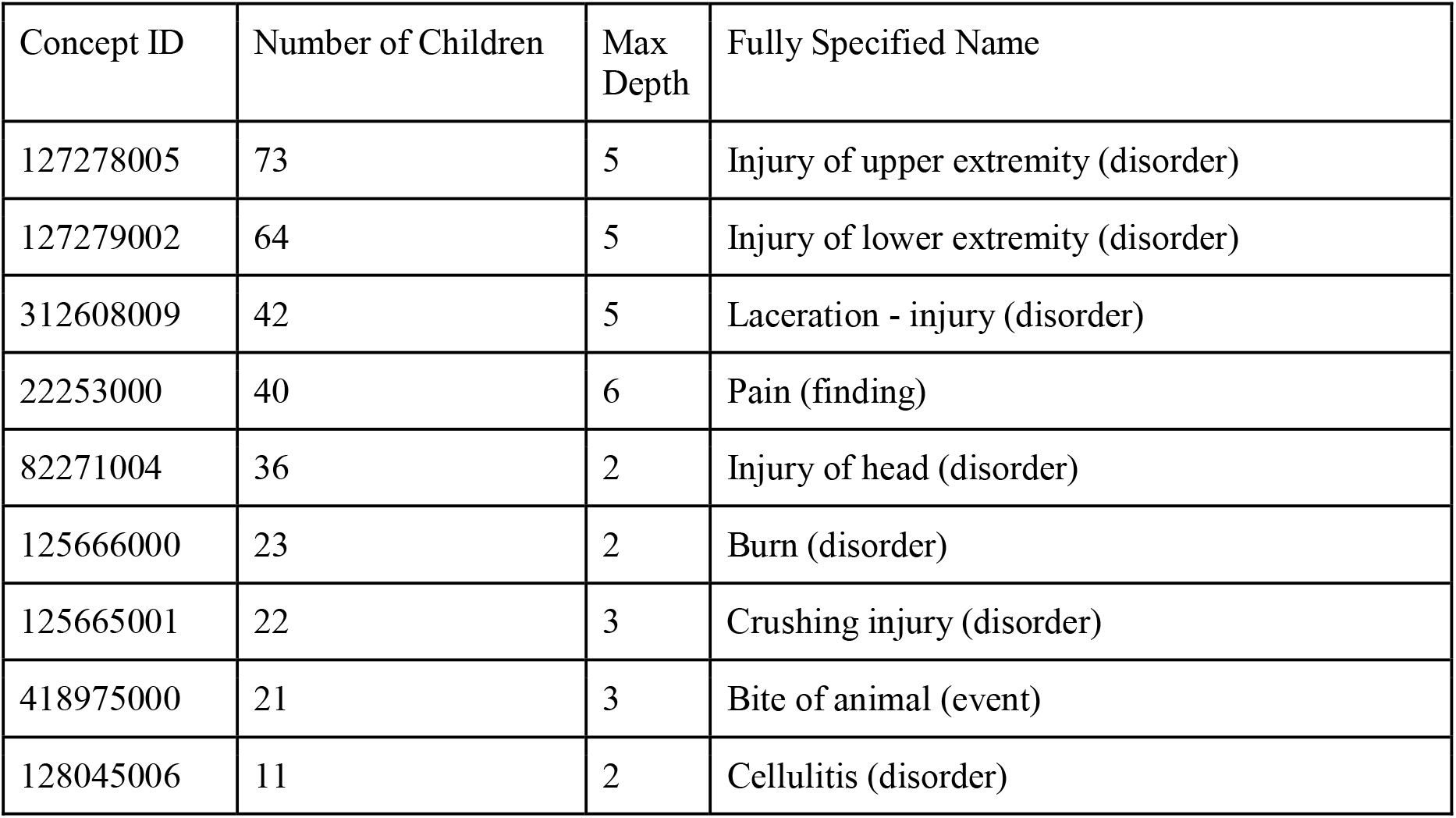
Top Level Concepts, Top 10 Ordered by Number of Children.

| Concept ID | Number of Children | Max Depth | Fully Specified Name |
| --- | --- | --- | --- |
| 127278005 | 73 | 5 | Injury of upper extremity (disorder) |
| 127279002 | 64 | 5 | Injury of lower extremity (disorder) |
| 312608009 | 42 | 5 | Laceration - injury (disorder) |
| 22253000 | 40 | 6 | Pain (finding) |
| 82271004 | 36 | 2 | Injury of head (disorder) |
| 125666000 | 23 | 2 | Burn (disorder) |
| 125665001 | 22 | 3 | Crushing injury (disorder) |
| 418975000 | 21 | 3 | Bite of animal (event) |
| 128045006 | 11 | 2 | Cellulitis (disorder) |

### 4.3 Error analysis

1,086 entries failed to match. We randomly selected 121 entries (10%) for further analysis. We manually reviewed each of these entries and then normalized them by removing ambiguous abbreviations (e.g., “BRADY” -> “Bradycardia”), punctuation (e.g, “\NUMBNESS” -> “Numbness”), or sentence structure (e.g, “FOR EVAL/? SZ”-> “Seizure”) that may have been present. Using a process similar to that described by Zhou^29^, we then manually mapped each normalized term to their corresponding SNOMED CT concept if available (**Table 8**). Post-normalization, each complaint was scored as an exact match (e.g., aortic stenosis->aortic stenosis) a partial match (e.g., frontal lobe mass -> brain mass) or, if no match could be found in SNOMED CT, as missing. We then examined why entries could not be matched (**Table 9**).

**Table 8:** Error Analysis.

|  | # SNOMED CT<br>(n=120) |
| --- | --- |
| Exact | 57 |
| Partial | 58 |
| Missing | 5 |

\*An exact match denotes a PP that can be mapped to a single SNOMED concept that accurately captures the scope and granularity of the PP entered at triage. A partial match means that the PP entered is similar to, but distinct from, existing SNOMED phraseology.

**Table 9:** Error Analysis.

| **Ambiguous Terms** | **n=120 (%)** | **Examples** | **Matching Concept** |
| --- | --- | --- | --- |
| Lexical Variant | 41 (34%) | Febrile | Fever (finding) |
| Logical Operator | 4 (3%) | S/P Bee Sting | Bee sting (disorder) |
| Multi-complaint | 13 (11%) | SI/SA | Suicidal (finding)<br>Suicide attempt (event) |
| Abbreviation | 11 (9%) | FB | Foreign body (disorder) |
| Malformed | 5 (4%) | BICYCLIST ST5RUCK | Bicycle accident (event) |
| Granularity | 29 (24%) | T9 MASS | Mass of thoracic vertebrae (finding) |
| Exact match | 17 (14%) |  |  |

### 4.4 Proposed SNOMED Additions

We discovered a set of 18 concepts that were not yet in SNOMED that we believe should be added to future revisions of SNOMED (**Table 10**). Three of these concepts have since been added to SNOMED in the 2018 release. We will request inclusion of these terms via the US Edition SNOMED CT Content Request System.

**Table 10:**
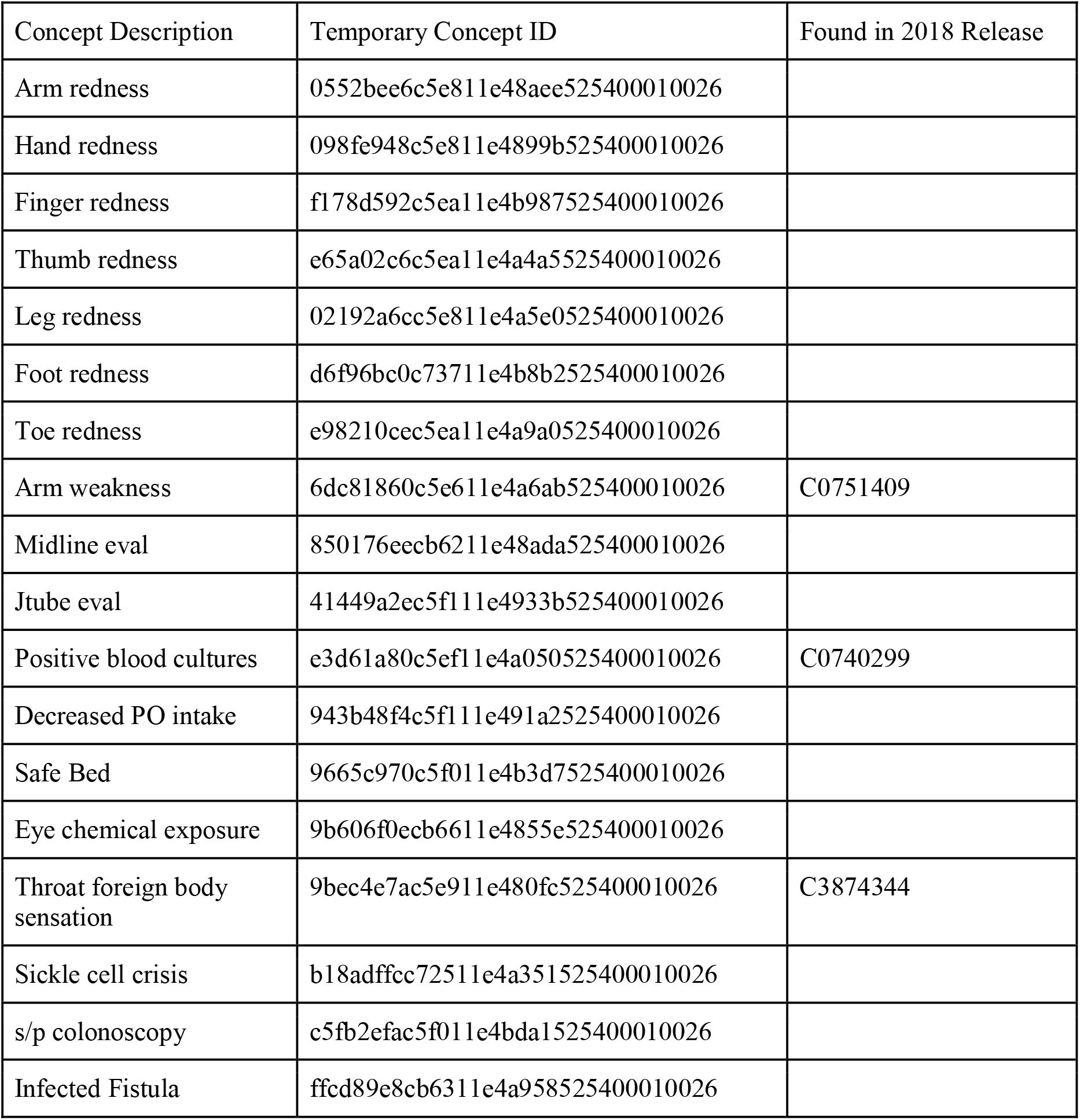
Terms that need to be added to SNOMED CT.

## 5. DISCUSSION

### 5.1 Pre/Post coordination

We examined the balance between pre-coordination and post-coordination and its implications at the point of entry.

With pre-coordination, a concept can be represented using one single concept identifier while post-coordination consists of two or more concepts that are used to represent a single complaint^30^. For example, entering the complaint of “left sided chest pain” in a post-coordinated fashion entails the entry of three concepts starting with “pain”, adding the location modifier “chest” and then a laterality modifier “left”. Alternatively, entry of this concept using pre-coordination would require selection of just one concept “left side chest pain”. We are not aware of any EDIS that supports post-coordination of presenting problems. Since our initial goal was to create an ontology that could be used in existing EHR’s (Goal 2), post-coordination for anatomical location would be impractical. Furthermore, post-coordination could result in clinically nonsensical concepts, concept duplication, and inefficiency of concept composition.^26^ Therefore, we chose to use pre-coordinated SNOMED CT terms, using post-coordination only to denote laterality. Though laterality may not be important to other users of SNOMED CT, laterality is extremely important in emergency departments. Wrong side errors is a well recognized problem in medicine. For example, a patient with right wrist pain is more likely to get the correct x-ray than a patient with a presenting problem of wrist pain.

### 5.2 Polyhierarchy

We modeled presenting problems using a hierarchical approach to improve the usability of the terminology in identifying complaints and their relationships to one another. Building off SNOMED CT, we were able to exploit the hierarchy and relationships that have already been established to streamline our implementation.

Structural representation of concepts within the terminology is important both for finding concepts and understanding relationships between concepts. Terminologies are typically structured in either a flat (i.e., list) form or in a hierarchical form. Flat terminologies provide no relationship information connecting related concepts. As such, they are of limited utility when searching for concepts that are associated. Conversely, in a hierarchical structure, relationships between concepts are clearly defined and can be used computationally to create complex queries.

Most ontologies defined for biomedicine use a Basic Formal Ontology^31^ (BFO) approach where a monohierarchy is asserted. In a monohierarchy, each concept has exactly one parent, while in a polyhierarchy, each child may have more than one parent. In a monohierarchy, the concept ‘Right lower quadrant pain’ would be required to have a parent of *either* ‘Right sided abdominal pain’ *or* ‘ Lower abdominal pain’, but not both. Coercing ‘RLQ pain’ into one of these categories would force the ontology to diverge from a clinicans’ mental model of the presenting problem. The prior probability of disease differs greatly given the location of pain in a patient’s abdomen. Therefore, specifying the exact location of abdominal pain is critical. In fact, we noted that users would override the system and enter presenting problems not found in the ontology when an appropriately specific presenting problem was not found. Although this could be represented as pain, with location right lower quadrant, which is part of the abdomen through post-coordination, most EHR’s do not support post-coordination. We discuss the tradeoffs of pre-coordination and post-coordination more extensively in the previous section. Because of the aforementioned reasons and current limitations of EHR’s, we felt strongly that only a polyhierarchal ontology would suit our original goals.

Existing standards, such as International Statistical Classification of Diseases and Related Health Problems (ICD) and Current Procedural Terminology (CPT) codes, are well suited for public health and billing applications but use monohierarchy; this limits the granularity of a concept as a term can only ever have one parent. Therefore, ICD10 was specifically rejected as the host terminology because it is not polyhierarchical.

Polyhierarchical structures mirror clinical reasoning and, as our ontology expands, provide a framework to add new concepts. In contrast, flat terminologies provide no relationship information connecting various concepts and invariably fail to provide the level of granularity that maximizes utility.

### 5.3 Data Mapping

Terms were mapped to a reference terminology, SNOMED CT, where they can be described using formal relationships or descriptions (e.g., chest pain “is a” disorder that has a “finding site” of “chest”).^26^ SNOMED CT is particularly useful for emergency department presenting problems, where the patient’s complaint may be a symptom, the name of a disease, a physical finding, or an event.

Owing to its widespread use and robust nomenclature SNOMED has become the de facto standard for clinical terminology. Mapping to SNOMED enhances data re-use and facilitates translation into non-english languages.

### 5.4 Comparative Analysis

There have been various attempts to create coded lists of chief complaints over the last 4 decades. (**Table 11**) shows comparison information of openly available and published PP lists as well as the PP list described here. Most notable differences are the polyhierarchical structure and mapping to SNOMED CT.

**Table 11:**
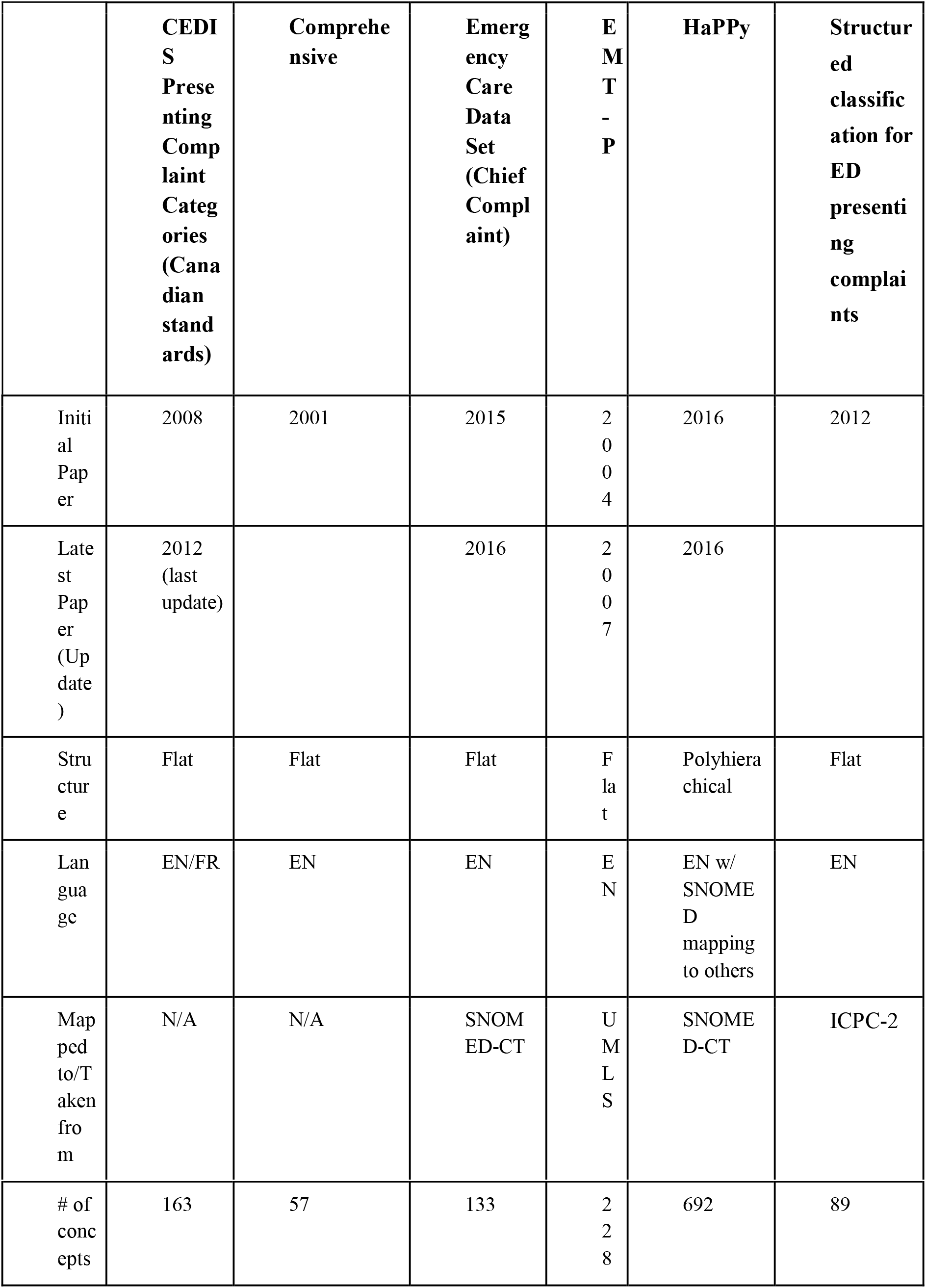
Comparative Analysis of Presenting Problem Approaches.

### 5.5 Limitations

Although we had a multi-center informatics team, our interface terminology was generated from presenting problems obtained from one tertiary academic medical center. This terminology may not be generalizable to other emergency departments practicing in another geographical areas or with a different patient population. Extensive external validation across different care settings (urban, rural, etc.) will be needed to further refine this interface terminology.

A major limitation of this work is that it only represents adult presenting problems, not pediatric presenting problems in the emergency department. Pediatric presenting problems were deliberately excluded as our data did not include pediatric patients.

Terms not present in SNOMED CT will require new concepts to be added and the appropriate mappings to be created. Terminologists at SNOMED CT may differ in their mapping of concepts. Although our terminology adheres to Rosenbloom et al.’s criteria for an interface terminology, the presenting problems were generated in an empirical fashion and the balance between pre- and post-coordination was determined by content experts.

Lastly, we recognize that all classification schemes are a product of expert opinion. Like any classification system, ours could unwittingly be manipulated to over or under represent certain situations, events, and conditions. For example, both ‘snake bite’ and ‘animal bite’ appear in our ontology. Had ‘snake bite’ been omitted, it could have falsely underestimated the incidence of these events, which could inadvertently lead to reduced funding for research and antidote development. To help mitigate these challenges we intend to regularly update and refine our ontology based on community feedback.

### 5.6 Implementation Suggestions

Whereas our ontology will be updated on an ongoing basis, we recommend that developers store PPs as the text entered by the user during the patient visit, as opposed to translating the user’s input into a SNOMED code. This ‘late binding’ will allow for retroactive reclassification of complaints as our ontology is refined and matures.

Similarly, we recommend a user interface design that permits concurrent autocompleted items from our ontology to be used alone or in combination with free text entry from the provider.^32^ This hybrid approach will increase the generation of structured data while still enabling providers to enter free text information that reflects their clinical judgment. These free text additions should be captured and submitted as candidates for future inclusion in our ontology

### 5.7 Future Directions

The next steps are to deploy this ontology to additional institutions, both retrospectively and prospectively. In the retrospective arm, we would see how well our interface terminology maps previously documented presenting problems. In the prospective arm, we would deploy the ontology, and after a washout period, we would analyze how well the ontology captures presenting problems. We would then analyze usage for any terms that did not match, and consider adding them to the ontology, using the same heuristics developed earlier. This could mean either adding new interface terminology, or new concepts. We will continue this iterative development cycle until we reach saturation of concepts. An important future direction for any ontology is securing organizational and financial support for its ongoing maintenance and curation. Part of this maintenance would be adding the ontology to the National Library of Medicine’s Value Set Authority Center (VSAC).^33^ As part of our post-publication evaluation process, we will require users to agree to an annual survey, so that we can better understand how the ontology is being used as well as penetrance.

Another exciting future direction would be to use machine learning to help curate the ontology by suggesting new concepts and interface terminology. We have previously developed machine learning models to predict presenting problems based on triage data. We already use this to help users input presenting problems.^32^ We could similarly use machine learning to help curate the ontology.

We also plan to work with the relevant national disease registries, quality registries^34^, research networks^35^, EHR vendors, and standards organizations to further refine and develop this ontology.

As existing ontologies for presenting problems (chief complaints) already exist, it would be interesting to develop a cross-walk to map this ontology to prior ontologies, as not all ontologies map to a standard ontology such as SNOMED CT.

Lastly, we believe this reference set could be adapted to any language. A possible future direction would be to create an interface terminology for other English language regions such as the United Kingdom or Australia, followed by other languages such as Spanish or French. This would be particularly

## 6. CONCLUSIONS

We present the HierArchical Presenting Problem ontologY (HaPPy). This ontology was empirically derived then iteratively validated by an expert consensus panel. HaPPy contains 692 presenting problem concepts, each concept being mapped to SNOMED CT. This freely sharable ontology should help to facilitate presenting problem based quality metrics, research, and patient care.

## 7. CLINICAL RELEVANCE STATEMENT

Accurately capturing presenting problems is a vital tool for understanding patterns of patient visits, clinical decision support, quality measures, syndromic surveillance, and research. Our empirically derived, expert validated system provides the first freely sharable, robust ontology capable of accomplishing this task.

## 8. ACKNOWLEDGEMENTS

Administrative support was partially funded by an American College of Emergency Physicians Section Grant.

We would like to acknowledge Stacie Jones for administrative support, as well as Laura Heermann Langford, Kevin Coonan and Adam Landman for their participation in the consensus process.

## 10. HUMAN SUBJECTS PROTECTIONS

This project was reviewed by the Committee on Clinical Investigations at Beth Israel Deaconess Medical Center and a determination (#2019D000313) was made that this activity did not constitute Human Subjects Research and no further review was required.

## 11. DATA AVAILABILITY

As a derivative work of SNOMED CT, The HierArchical Presenting Problem ontologY (HaPPy) is released freely to anyone with a valid Snomed CT license. The ontology can be downloaded via GitHub at https://github.com/hornste/happy

