## Supplemental Tables for "Consensus Development of a Modern Ontology of Emergency Department Presenting Problems – the HierArchical Presenting Problem ontologY (HaPPy)"

**Supplementary Table 1: Synthesis of lexical variants**

| Left | L, Lt, (L) | side, -side, sided, -sided |
| --- | --- | --- |
| Right | R, Rt, (R) | side, -side, sided, -sided |
| Bilateral | B, Bilat, Bil, (B) |  |
| Upper Extremity | UE |  |
| Lower Extremity | LE |  |
| Fracture | Fx |  |
| Injury | Inj |  |
| Abnormal | Abnl, Abn |  |
| Laceration | Lac |  |
| Evaluation | Eval |  |

**Supplementary Table 2**: Distribution of Concept Types and Their Usage

| **SNOMED Semantic Tag** | **No. of Concepts (n=457)** | **No. of Uses (n=112,986)** |
| --- | --- | --- |
| Disorder | 181 (40%) | 20,527 (18%) |
| Event | 12 (3%) | 2,516 (2%) |
| Finding | 228 (50%) | 85,681 (76%) |
| Procedure | 9 (2%) | 1,604 (1%) |
| Other | 27 (6%) | 2,658 (2%) |

**Supplementary Table 3: Concept Matching to SNOMED CT**

|  | **# SNOMED CT (n=168)** | **% SNOMED CT** |
| --- | --- | --- |
| Exact | 120 | 71% |
| Partial | 23 | 14% |
| Missing | 25 | 15% |
| *An exact match denotes a PP that can be mapped to a single SNOMED concept that accurately captures the scope and granularity of the PP entered at triage. A partial match means that the PP entered is similar to, but distinct from, existing SNOMED phraseology. | | |

**Supplementary Table 4: Comparative Analysis of Presenting Problem Approaches**

|  | **CEDIS Presenting Complaint Categories (Canadian standards)** | **Comprehensive** | **Emergency Care Data Set (Chief Complaint)** | **EMT-P** | **[XXX]** | **Structured classification for ED presenting complaints** |
| --- | --- | --- | --- | --- | --- | --- |
| Initial Paper | 2008 | 2001 | 2015 | 2004 | 2016 | 2012 |
| Latest Paper (Update) | 2012 (last update) |  | 2016 | 2007 | 2016 |  |
| Structure | Flat | Flat | Flat | Flat | Polyhierachical | Flat |
| Language | EN/FR | EN | EN | EN | EN w/ SNOMED mapping to others | EN |
| Mapped to/Taken from | N/A | N/A | SNOMED-CT | UMLS | SNOMED-CT | ICPC-2 |
| # of concepts | 163 | 57 | 133 | 228 | 690 | 89 |
